# Identifiers for the 21st century: How to design, provision, and reuse persistent identifiers to maximize utility and impact of life science data

**DOI:** 10.1101/117812

**Authors:** Julie A McMurry, Nick Juty, Niklas Blomberg, Tony Burdett, Tom Conlin, Nathalie Conte, Mélanie Courtot, John Deck, Michel Dumontier, Donal K Fellows, Alejandra Gonzalez-Beltran, Philipp Gormanns, Jeffrey Grethe, Janna Hastings, Henning Hermjakob, Jean-Karim Hériché, Jon C Ison, Rafael C Jimenez, Simon Jupp, John Kunze, Camille Laibe, Nicolas Le Novère, James Malone, Maria Jesus Martin, Johanna R McEntyre, Chris Morris, Juha Muilu, Wolfgang Müller, Philippe Rocca-Serra, Susanna-Assunta Sansone, Murat Sariyar, Jacky L Snoep, Natalie J Stanford, Stian Soiland-Reyes, Neil Swainston, Nicole Washington, Alan R Williams, Sarala Wimalaratne, Lilly Winfree, Katherine Wolstencroft, Carole Goble, Christopher J Mungall, Melissa A Haendel, Helen Parkinson

## Abstract

In many disciplines, data is highly decentralized across thousands of online databases (repositories, registries, and knowledgebases). Wringing value from such databases depends on the discipline of data science and on the humble bricks and mortar that make integration possible; identifiers are a core component of this integration infrastructure. Drawing on our experience and on work by other groups, we outline ten lessons we have learned about the identifier qualities and best practices that facilitate large-scale data integration. Specifically, we propose actions that identifier practitioners (database providers) should take in the design, provision and reuse of identifiers; we also outline important considerations for those referencing identifiers in various circumstances, including by authors and data generators. While the importance and relevance of each lesson will vary by context, there is a need for increased awareness about how to avoid and manage common identifier problems, especially those related to persistence and web-accessibility/resolvability. We focus strongly on web-based identifiers in the life sciences; however, the principles are broadly relevant to other disciplines.

## Introduction

The issue is as old as scholarship itself: readers have always required persistent identifiers in order to efficiently and reliably retrieve cited works. ‘Desultory citation practices’ have been thwarting scholarship for millennia [1]. While the Internet has revolutionized the *efficiency* of retrieving sources, the same can not be said for reliability: it is well established that a significant percentage of cited web addresses go “dead” [2]. This process is commonly referred to as link rot because availability of cited works decays with time [3,4]. Although link rot threatens to erode the utility and reproducibility of scholarship [5], it is not inevitable: link persistence has been the recognized solution since the dawn of the Internet [6]. However, this problem, as we will discuss, is not at all limited to referencing journal articles. The life sciences have changed a lot over the past decade as the data have evolved to be ever larger, more distributed, more interdependent, and more natively web-based. This transformation has fundamentally altered what it even means to ‘reference’ a resource; it has diversified both the actors doing the referencing and the entities being referenced. Moreover, the challenges are compounded by a lack of shared terminology about what an ‘identifier’ even is. Box 1 delineates the key components of an identifier used throughout this paper; all technical terms are in fixed width font and defined in the glossary.

### Box 1. Anatomy of a persistent identifier

An **identifier** is a sequence of characters that identifies an entity. The term ‘**persistent identifier**’ is usually used in the context of digital objects that are accessible over the Internet. Typically, such an identifier is not only persistent but also actionable [7]: it is a **Uniform Resource Identifier (URI)** [8], usually of type http/s, that you can paste in a web browser address bar and be taken to the identified source.

An example of an exemplary **URI** is below; it is comprised of ASCII characters and follow a pattern that starts with a fixed set of characters (URI pattern). That URI pattern is followed by a **Local ID** -- an identifier which, by itself, is only guaranteed to be locally unique within the database or source. A local ID is sometimes referred to as an ‘accession’.

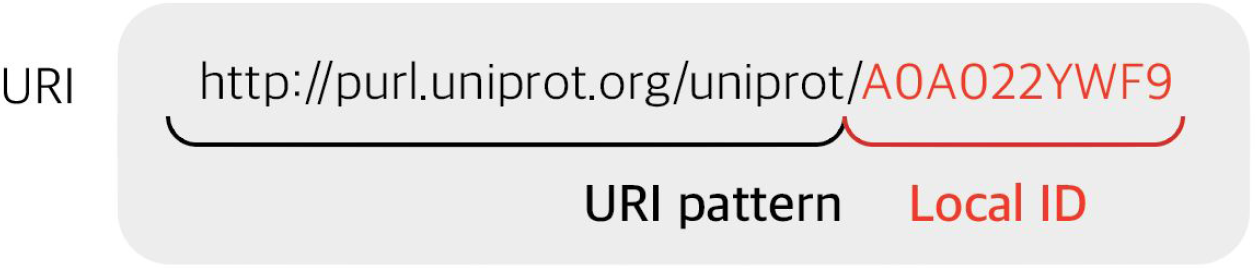

Formally breaking down a URI into into these two components (URI pattern and local ID) makes it possible for meta resolvers to ‘resolve’ entities to their source. This practice also facilitates representation of a URI as a **compact URI (CURIE)**, an identifier comprised of <**Prefix**>:<**Local ID**> wherein **prefix** is deterministically convertible to **a URI pattern** and vice-versa. For instance, the above URI could be represented as uniprot:A0A022YWF9. This deterministic conversion makes it easy for meta resolvers as well, e.g., http://identifiers.org/uniprot:A0A022YWF9.

Suboptimal identifier practice is artificially constraining what can and cannot be done with the underlying data: it not only hampers adherence to FAIR principles (findability, accessibility, interoperability, and reuse) [9], but also compromises mechanisms for credit and attribution. This article seeks to provide pragmatic guidance and examples for how actors in life science research lifecycle should handle identifiers. Optimizing web-based persistent identifiers is harder than it appears; there are a number of approaches that may be used for this purpose, but no single one is perfect: Identifiers are reused in different ways for different reasons, by different consumers. Moreover, digital entities (e.g., files, such as an article), physical entities (e.g., tissue specimens), living entities (e.g. Dolly the sheep), and descriptive entities (e.g., ‘mitosis’) have different requirements for identifiers [10].

The problem of identifier management is hardly unique to the life sciences; it afflicts every discipline from astronomy [3] to law [11]. Towards this end, several groups (**Supplemental Text S1**) have been converging on identifier standards that are broadly applicable [9, 12-14]. Building on these efforts and drawing on our experience in integrating and accessing data from a large number of sources, we outline the identifier qualities and best practices we consider particularly important in the context of large-scale data integration in the life sciences. In **Lessons 1-9** (**Table 1**) we propose actions for data providers when designing new identifiers, maintaining existing identifiers, as well as when reusing and referencing identifiers from other datasets. In **Lesson 10**, we conclude with guidance for data integrators and redistributors on how best to reference multiple identifiers from diverse sources. More often than not, life science data providers often invent or organically grow their own identifier systems without a firm grasp of the lasting implications. Data providers are urged to take a long-term view of the scope and lifecycle of data and the identifiers that they issue, and to consider using existing identifier platforms and services [13] where appropriate.

**Table 1.**
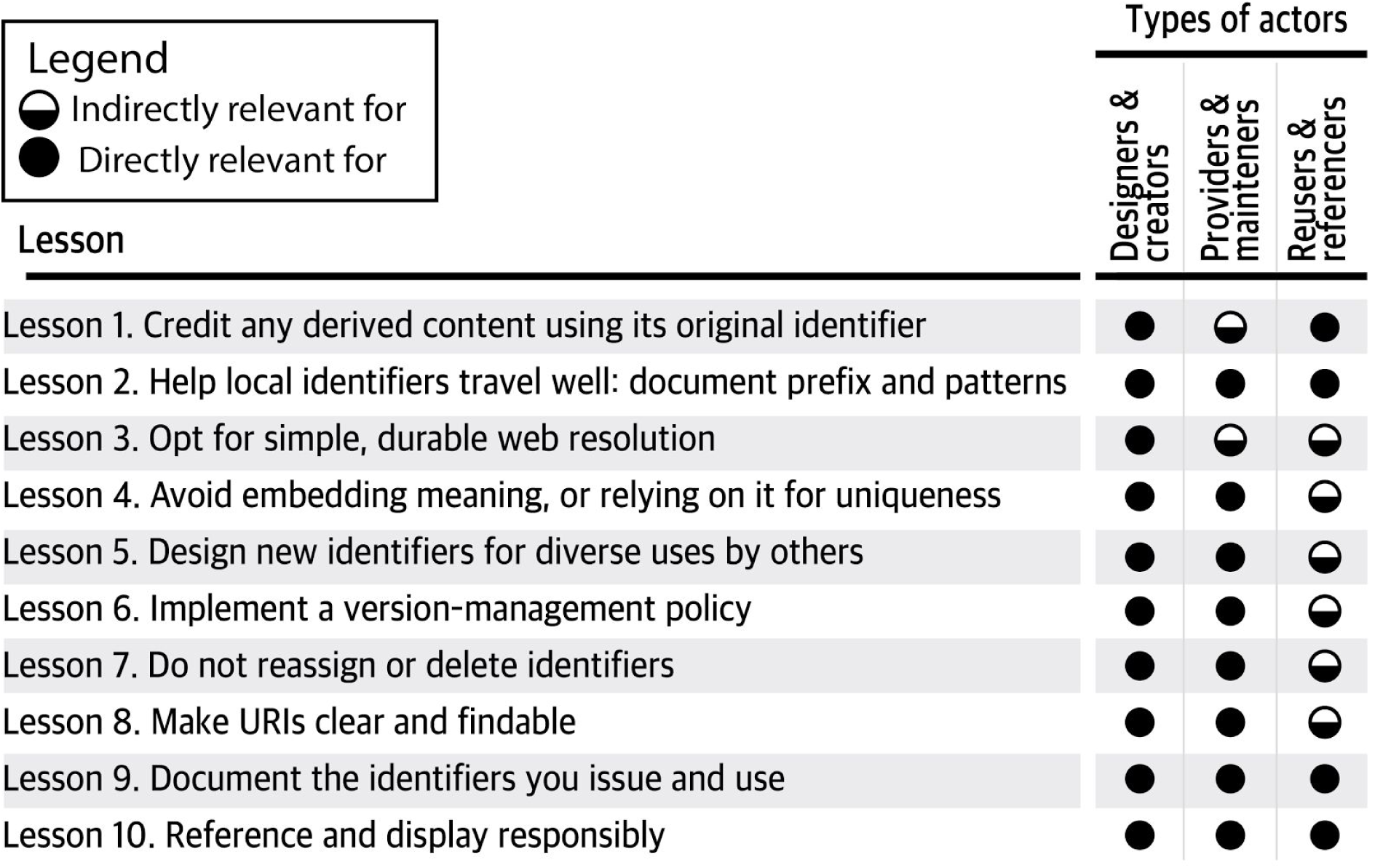
A summary of the 10 recommendations, their direct or indirect impact on different kinds of identifier actions.

Throughout this document, the word “must” is reserved for practices that ensure against the collision, ambiguity, or inaccessibility of items referenced by identifiers; instances of “must” are also often specific to particular design choices. We use the word “should” to convey that the tradeoffs must be understood and carefully weighed before choosing a different course (eg. consistent with IETF RFC2119 [15]). Terms that appear in fixed-width font are defined in the supplemental glossary (**Supplemental Table S2**).

There is no one in science that is unaffected by identifiers. **Table 1** details three basic roles one might play in the scholarly landscape and how identifiers are relevant in these contexts. Who are designers and creators? These are databases, but also those that submit supplemental data to archives, and anyone creating structured data. Who are the providers and maintainers? These are databases as well, but also services and indices that support web resolution and data validation. Who are the reusers and referencers? These are the “Research Data Parasites” [16], but also your average author: while authors may specify an identifier for a resource (e.g. a gene or antibody), more often identifiers are contextually inferred by the journals or curators, whether prey- or post-publication.

Many of the following recommendations are applicable during the planning and identifier conceptualization phase, i.e. before any identifiers are created. The retrofitting (especially Lessons 1, 4, 5, and 6) of existing identifiers can sometimes be too difficult or may even make matters worse: for instance changing existing identifiers introduces the need for systems that can recognize the variations for what they are; such overhead can outweigh potential benefits. Each of the lessons is relevant to the basic classes of identifier actions (design, provision, reuse **Table 1**) within the ecosystem of data providers and integrators. These actions in turn are relevant to anyone on the spectrum of seven basic roles ranging from those that publish their own data to those that provide applications on top of others’ data (**Figure 1**). Even if we largely agree on what makes for a good persistent identifier (**Table 2**), actual implementation often falls short. No provider is perfect and no two are alike, hence the objective is to learn from each other’s diverse experiences. All of the negative examples herein are anonymized variations of real-world identifiers that we have had to work with.

**Figure 1.**
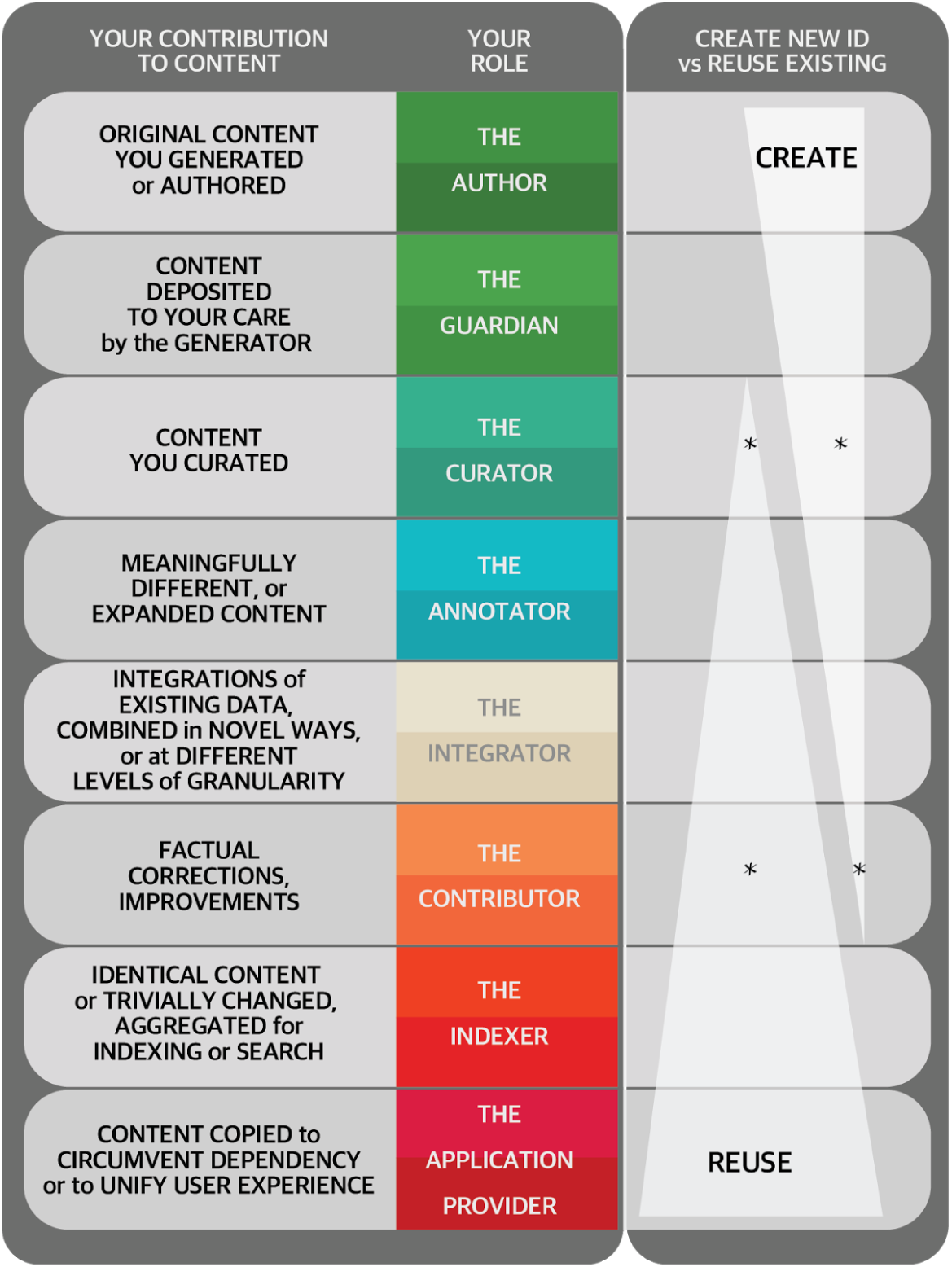
Contributions and roles related to content as they correspond to identifier creation vs reuse. The decision about whether to create a new identifier, or reuse an existing one depends on the role you play in the creation, editing, and republishing of content; for certain roles (and when several roles apply) that decision is a judgement call. Asterisks convey cases in which the best course of action is often to correct/improve the original record in collaboration with the original source; the guidance about ID creation versus reuse is meant to apply only when such collaboration is **not** practicable (and an alternate record is created). It is common that a given actor may have multiple roles along this spectrum; for instance, a given record in monarchinitiative.org may reflect a combination of a) corrections Monarch staff made in collaboration with the original data source, b) post-ingest curation by Monarch staff, b) expanded content integrated from multiple sources.

**Table 2.**
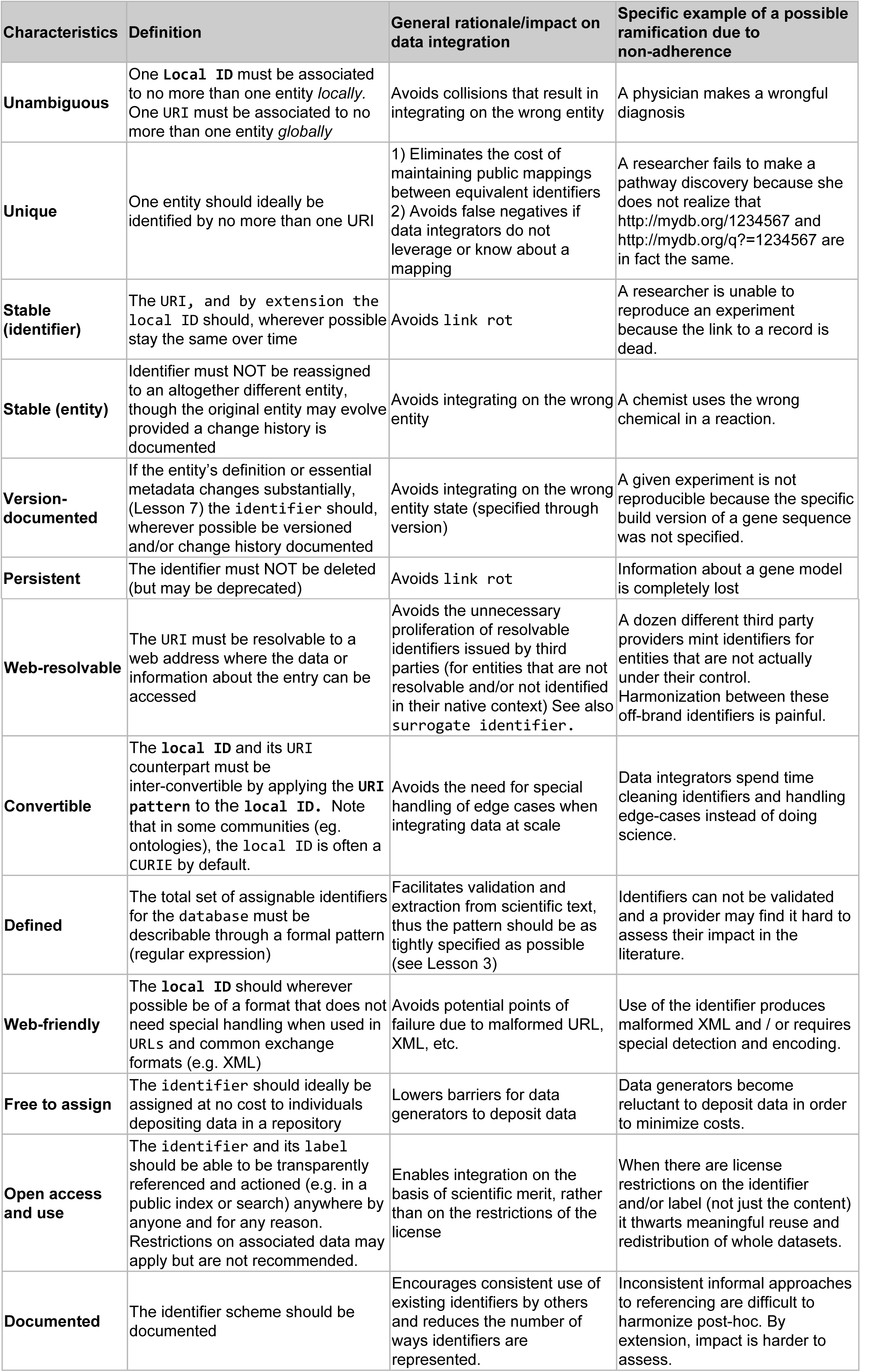
Desirable characteristics for database identifiers in the life sciences.

### Lesson 1. Credit any derived content using its original identifier

If you manage an online database (repository, registry, or knowledgebase), consider its role in identifying and referencing the knowledge that it publishes. We advise that you only create your own identifiers for new knowledge (**Figure 1**). Wherever you are referring to existing knowledge, do so using existing identifiers (Lesson 10): otherwise, wherever the 1:1 relationship of identifier:entity breaks down, costly mapping problems arise. Whether or not you create a new ID, it is vital to credit any derived content using its indigenous identifiers [10]; to facilitate data integration, all such identifiers should be machine processable and transparently mapped.

### Lesson 2. Help local identifiers travel well: document prefix and patterns

If you reference others’ data, or anticipate your data *being referenced by others,* consider how you document your identifiers. Note that you may not know *a priori* how your data may be used. Data does not thrive in silos: it is most useful when reused, broken into parts and integrated with other data, for instance in database cross references (“db xrefs”). In spite of how important identifiers are to this process, the confusion with identifiers often starts with the basics, including what the “identifier” even is. A local ID (**Box 1**) is an identifier guaranteed only to be unique in a given local context (eg. a single provider, a single collection, etc.), and sometimes only within a specific version; as such, it is poorly suited to facilitate data integration because it can collide when considered in a more global landscape of many such identifiers. For instance, the local ID “9606” corresponds to numerous entities whose local accessions are based on simple digits, including: a <u>Pubmed article</u>, a <u>CGNC gene</u>, a <u>PubChem chemical</u>, as well as an <u>NCBI taxon</u>, a <u>BOLD taxon</u>, and a <u>GRIN taxon</u>. Local IDs therefore need to be contextualized in order to be understood and accessed (resolved) on the web. This is often accomplished through the use of a prefix, which should be documented. If this is overwhelming, don’t forget that there are meta resolvers and services built to help for exactly this reason (see **Lesson 3**).

Uniform Resource Identifiers (URIs) are identifiers that resolve on the web. “Cool URIs don’t change” [6] because when, they *do change* (or disappear) all existing references break. In the context of academia alone, “reference rot” problem impacts one in five publications [4]. Despite link rot vulnerability, the global http/s URI (**Box 1**) is the best available identifier form for machine-driven global data integration because a) the http URI is a widely adopted IETF standard and b) the http URI’s uniqueness is ensured by a single well-established name-granting process (<u>DNS</u>). However, the length of URIs can make them unwieldy for tasks involving human readability even within structured machine-parsable documents. Compact URIs (CURIEs [17], **Box 1**) are a mature W3C standard that is well established in some contexts (e.g. JSON-LD and RDFa) as they enable URIs to be understood and conveniently expressed. We the authors are not absolutist about anyone using CURIEs; however, we agree that the features that make for good URIs also happen to make CURIEs possible (for those who wish to use them) (Supplementary Text S3).

Thus if you are a database provider, it is in your best interests to document and preferably register a) the prefix (**Box 1**) that you would like others to use and b) its binding to a URI pattern (**Box 1**). Your chosen prefix should be unique, at least among datasets that are likely to be used in the same context. Supplementary Table S4 contains a list of registries that may be suitable depending on the kind of data. PrefixCommons [18] is a platform designed to enable such registries to make more informed decisions about which prefixes to issue and utilized and for any given integrator to publish the mappings that they happen to use.

### Lesson 3. Opt for simple, durable web resolution

A core component of persistent identification is redirection, the absence of which makes it extremely difficult to provide stable identifiers. When designing (or refining) your http URI strategy:

- **Consider a resolution provider before doing it yourself.** If you are a database provider, you must implement an http URI pattern (**Figure 1 panel B**) for **local IDs** to be resolvable to a web page. If you choose to outsource to a resolver service, use an approach that adheres to best practice [13] (e.g. DOI (<u>DataCite</u>, <u>CrossRef</u>), Identifiers.org, Handle.net, [PURL](https://purl.oclc.org/) (now via InternetArchive), [EPIC](http://www.pidconsortium.eu/), [ARK](https://wiki.ucop.edu/display/Curation/ARK)) and be mindful of your constraints regarding cost, metadata ownership, turnaround time, etc. (See **Supplemental Text S5** for a more comprehensive list of considerations.) Some of these resolver services can even provide content negotiation for different encodings of your data [13] and make it easier to provide direct access to data, metadata, and persistence statements [19]. If you have the resources to support your own persistent URIs, design these to be “cool” [6]; this is most easily achieved by keeping URIs simple.
- **Avoid inclusion of anything that is likely to change or lapse**, including administrative details (e.g. grant name) or implementation details such as file extensions (‘resource.html’), query strings (‘param=value’), and technology choices (‘.php’). Never embed the **local ID** in the query part of a URI eg. http://example.com/explore?record=A123456.
- **Omit trailing characters after the local ID.** In all cases, the URI pattern must include the protocol (e.g. https://) and, if applicable, trailing slash or other delimiters. Trailing characters after the local ID are discouraged as they unnecessarily increase the variability with which the identifier is represented and also complicate straightforward appending of the local ID (requiring that tokens such as $id hold the place of the **local ID** in the URI pattern eg http://example.com/$id/view.do).
- **Avoid unnecessary detail.** Detail in ‘persistent’ identifiers creates complexity that must be managed in perpetuity. Make every attempt to limit the degree of path nestedness (eg. do http://example.com/A123456 rather than http://example.com/vertebrates/mammals/rodents/rat/white-rat/A123456): see also Lesson 5 regarding types and meaning. The compact URI approach can work with any resolver(s): see for instance examples 4 and 5 in **Figure 2**. By choosing a single URI pattern, you make it possible for others to resolve your identifiers simply (**Figure 2 panel A**) without their having to know the type and its syntax in http URI. See also **Lesson 4** regarding omission of semantics.

Despite their differences, the examples in **Figure 2** share the most important features above.

**Figure 2.**
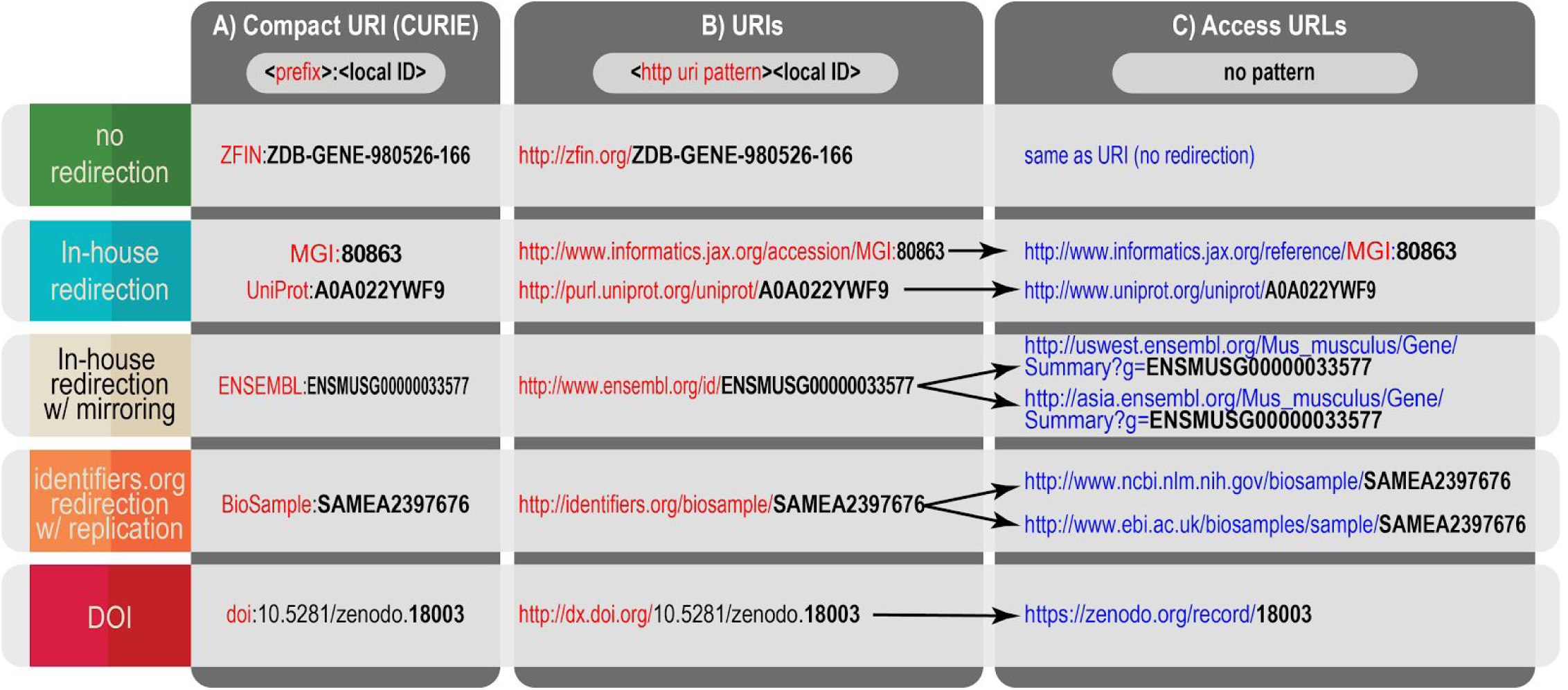
Examples of provisioning resolvable URIs: Compact URIs (CURIEs) (Panel A), URIs (Panel B) and Access URLs (Panel C) with no redirection (ZFIN), in house redirection (UniProt, and Ensembl), and 3rd party resolvers (using identifiers.org and DOI). In each case, the URI can be algorithmically derived from the CURIE because the **local ID** portion itself is included (unmodified) within the URI. Access URL design patterns differ substantially by provider and may change over time. As long as access URLs (and other ephemeral links) are not used as the referenced identifier, they can include prefix and colon (MGI) or not (Ensembl), they may include the entire local ID (Biosample) or not (DOI), and they may include type (MGI) or not (ZFIN).

### Lesson 4. Avoid embedding meaning, or relying on it for uniqueness

When designing new local IDs or http URIs, avoid embedding meaning or relying on it for uniqueness. The structure and scope of collections evolve, as does scientific understanding; minimizing the meaning embedded in identifiers makes them less vulnerable to obsoletion. In human genetics many genes were initially identified based on disease association; later the identification, nomenclature, and function of genes were separated into different activities. Meaning should only be embedded if it is indisputable, unchangeable and also useful to the data consumer (e.g. computer-processable). For instance, the type of entity imparts meaning to users and may fulfill these three criteria. When encountered, typing may be embedded, either within the **local ID** (ENS<u>MUSG</u>…), or within the http URI path (…/<u>gene</u>/12345), or both. In any case, if you opt to include type in the identifiers you issue, avoid relying on type for uniqueness: that is to say once a **local ID** eg. 12345 is assigned, it should never be recycled for another entity, even an entity of a different type for instance …/gene/12345 and …/patient/12345.

If you need the ability to convey meaning in a dense character space, you don’t need to do so in the identifier itself; consider instead implementing an entity label, for instance as is done in model organism nomenclature such as by Mouse Genome Informatics (label: [Kit^W^](http://www.informatics.jax.org/allele/MGI:1856232)/[Kit^W-v^](http://www.informatics.jax.org/allele/MGI:1856266), id: MGI:2171276). Labels are for human readability only; even if they are deemed durable, labels should not be treated as identifiers, nor should they appear within http URIs. URI patterns, if type-specific, require a corresponding type-specific prefix (e.g. for the Library of Integrated Network-based Cellular Signatures (LINCS), the prefix ‘LINCS-cell’ corresponds to http://lincs.hms.harvard.edu/db/cells/$id/ whereas the prefix ‘LINCS-protein’ corresponds to http://lincs.hms.harvard.edu/db/proteins/$id/). MGI implements both type-agnostic resolution (http://www.informatics.jax.org/accession/HGI:2442292) and type-specific destinations (http://www.informatics.jax.org/marker/MGI:2442292). Dual approaches like MGI’s can be helpful to different kinds of consumers: type-agnostic resolution is useful in cases such as data citation in the literature where a) the type of the identified entity is not of primary importance, or b) the type of the entity is already conveyed contextually, and/or c) where resolution is done systematically at scale and/or involves many and varied or volunteer contributors that may be difficult to coordinate. Type-specific resolution is useful in cases like bioinformatic research pipelines where embedded type may facilitate the human-led debugging process. If you support both kinds of resolution, it is best to document a) whether you intend for both to be treated as persistent b) what mapping support you provide.

Whether or not your URIs or your local IDs include type, you should provide other ways for humans and machines to determine the type of entity that is being identified; this is most often achieved via webservices (eg. **<u>as done via Monarch API</u>**), but ideally also within metadata landing pages [19,20] if provided.

### Lesson 5. Design new identifiers for diverse uses by others

Pre-existing identifiers should be referenced without modifications (see **Lesson 10**). However, if you create new **local IDS**, there are some design decisions that can facilitate their use in diverse contexts (spreadsheets, other databases, web applications, publications, etc.).

- **Avoid problematic characters.** Local IDs should, wherever possible comprise only letters, numbers and URL-safe delimiters. Omission of other special characters guards against corruption and mistranscription in many contexts; however, it is acceptable that the local ID be in CURIE format since modern browsers resolve colons without having to encode them. Although characters “/” and “?” are technically URL-safe, they are very problematic when used *within* the local ID as these characters are assumed to have special meaning and can complicate parsing of the identifiers, whatever forms they take. For the same reason, local IDs should ideally not contain ‘.’ except to denote version where appropriate (see **Lesson 7**).
- **Define a formal pattern and stick to it.** Local IDs must adhere to a formal pattern (regular expression); this facilitates the validation of URIs and improves the accuracy of mining identifiers from scientific text. Consider a fixed length of 8-16 characters (according to the anticipated number of required **local IDS**). A pattern may be extended if all available identifiers are issued, but existing identifiers should not be changed. To minimize **local ID** collisions at global scale, it is considerate to tightly specify your pattern (e.g. using one or more fixed letters). The regular expression should include a fixed, documented case convention. In most cases, it is advised that identifiers not rely on case for their uniqueness: if you assign ab-12345 to one entity and AB-12345 to *a different* entity, collisions due to mistranscription are more likely. Case-sensitive patterns are best reserved for when brevity is a constraint (e.g. millions of IDs are required and each ID has to be short enough to be printed on a vial label).
- **Avoid problematic patterns.** Consider using both letters and numbers in the local ID. This avoids misinterpretation as numeric data (e.g. truncation of leading zeros or conversion to exponents in spreadsheets). Some patterns can result in misinterpretation/corruption whether as dates (e.g. “may-15”), exponents (e.g. “5e1234”) [21], or as unintended words (e.g. “bad-12”). Such issues in gene names alone have been shown to impact 19% of life sciences papers [22]. A historically common, if thorny, identifier pattern is that ‘_’ and ‘:’ are often interconverted and it has come to be understood as compact notation, delimiting the prefix from the rest of the identifier. Therefore ‘_’ or ‘:’ should a) occur no more than once per identifier and b) should only be used if **local ID** are intended to be deterministically expanded to a resolvable http URI. For instance, if your intended prefix is ‘MyDB’, then either MyDB:gene-6622 or MyDB_gene-6622 are acceptable patterns, but MyDB_gene_6622 is problematic as it could result in three possible conversions by others, even if these are not intended: MyDB_gene:6622, MyDB:gene_6622, MyDB:gene:6622. Whatever pattern you adopt, document which variations you support resolution of, if any.

### Lesson 6. Implement a version-management policy

Whether you produce original data, or reference others’ data, consider the impact of changes. The nature, extent, and speed of data changes impact how data can be referenced and used. Document your chosen version management practice: If you issue identifiers, the change history for the entity should be either documented or queryable. Alternatively, the identifier itself can be versioned whether or not change history is also supported.

Explicit identifier versioning is recommended if the prevailing use of an *unversioned* identifier results in “breaking changes” (e.g., a change in the hypothesized cause of a disease). However, if new information about the entity emerges slowly and the changes are “non-breaking”, it is reasonable to instead maintain a machine-actionable change history wherein the changes are listed, and where they may also be categorized (eg. minor versus major changes). Versioning and change history work well together, especially when multiple types of changes overlap. Even where previous records are entirely removed, the URI should continue to resolve, but to a “tombstone” page (Lesson 7). A resource should communicate clearly what a version change refers to. UniProt and RefSeq use versions to reflect changes in sequence. Ensembl uses versions to reflect changes in sequence and splicing for transcript records but sequence alone for protein records. In each of these examples changes in annotation attached to a record does not alter the version.

There are two approaches to versioning, record-level and release-level; the latter is more common in the life sciences. Release-level versioning is usually performed for defined data releases. However, use cases vary; some user communities need to resolve individual archived entities via a deterministically-versioned URI pattern, for example as is done in Ensembl eg. http://e85.ensembl.org/id/ENSMUSG00000033577. In either case, whether or not you have the ability (or common use case) to maintain individually resolvable archived records, we strongly recommend supporting export to files so that users can archive the records they need. We also recommend making snapshots available for the database, whether in whole or in parts. [23]

If you version identifiers at the level of the individual record, you should version in the **local ID** after the ‘dot’, as per UniProt in **Table 3**; this provides continuity in your site and also enables a single prefix to be used with any version: UniProt:P12345.3 → http://www.uniprot.org/uniprot/P12345.3. If you do record-level versioning but dot suffixing is not practicable, we strongly recommend providing a transparent mapping between identifiers together with a mechanism for obtaining the latest version of the record (e.g. by inserting ‘/latest/’ in the URI path). Maintaining mappings between identifier versions without use of the dot is possible, but so difficult that few providers do it well. Other groups have discussed change management consideration and ‘content drift’ in more depth [2,24,25].

**Table 3.**
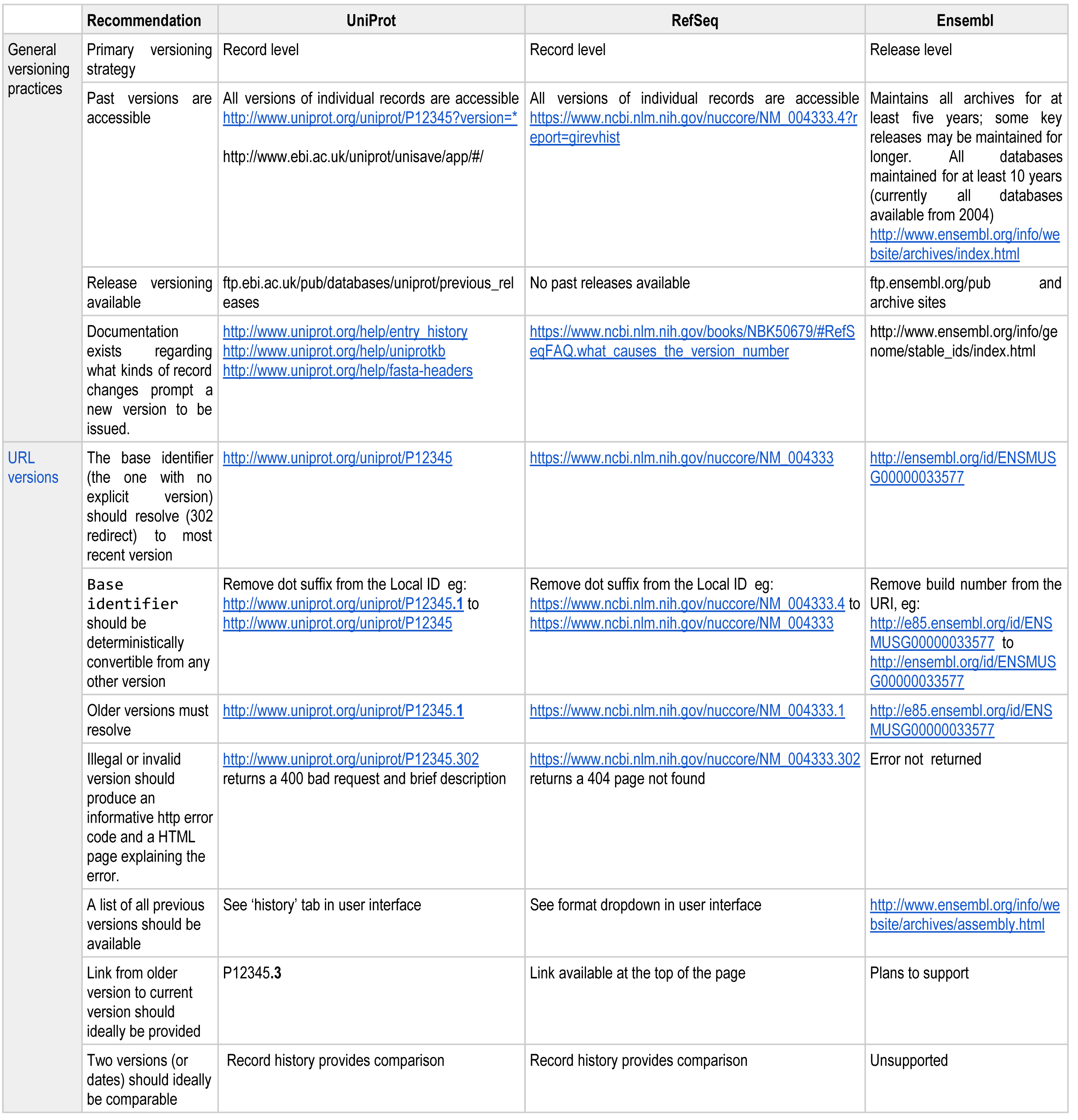
Recommendation for versioning.

### Lesson 7. Do not reassign or delete identifiers

Identifiers that you have exposed publicly, whether as http URIs or via APIs may be deprecated but must never be deleted or reassigned to another record. If you issue identifiers, consider their full lifecycle: there is a fundamental difference between identifiers which point to experimental datasets (GenBank/ENA/DDBJ, PRIDE, etc.) and identifiers which point to a current understanding of a biological concept (Ensembl Gene, UniProt record, etc.). While experimental records are less likely to change, concept descriptions may evolve rapidly; even the nature and number of the relevant metadata fields change over time. Moreover, the very notion of identity is often strongly impacted by relationships (e.g., between concepts or processes).

Extensive changes cannot be captured with numerical suffixing alone. For instance, taxonomists may split or merge species, pathologists may split or merge diseases, or hypothesized entities may be proven not to exist (e.g. vaccine-induced autism). Global initiatives (**Supplemental Text S1**) are actively exploring identifier strategies for such use cases. In the meantime, consider **Table 4** recommendations.

**Table 4.**
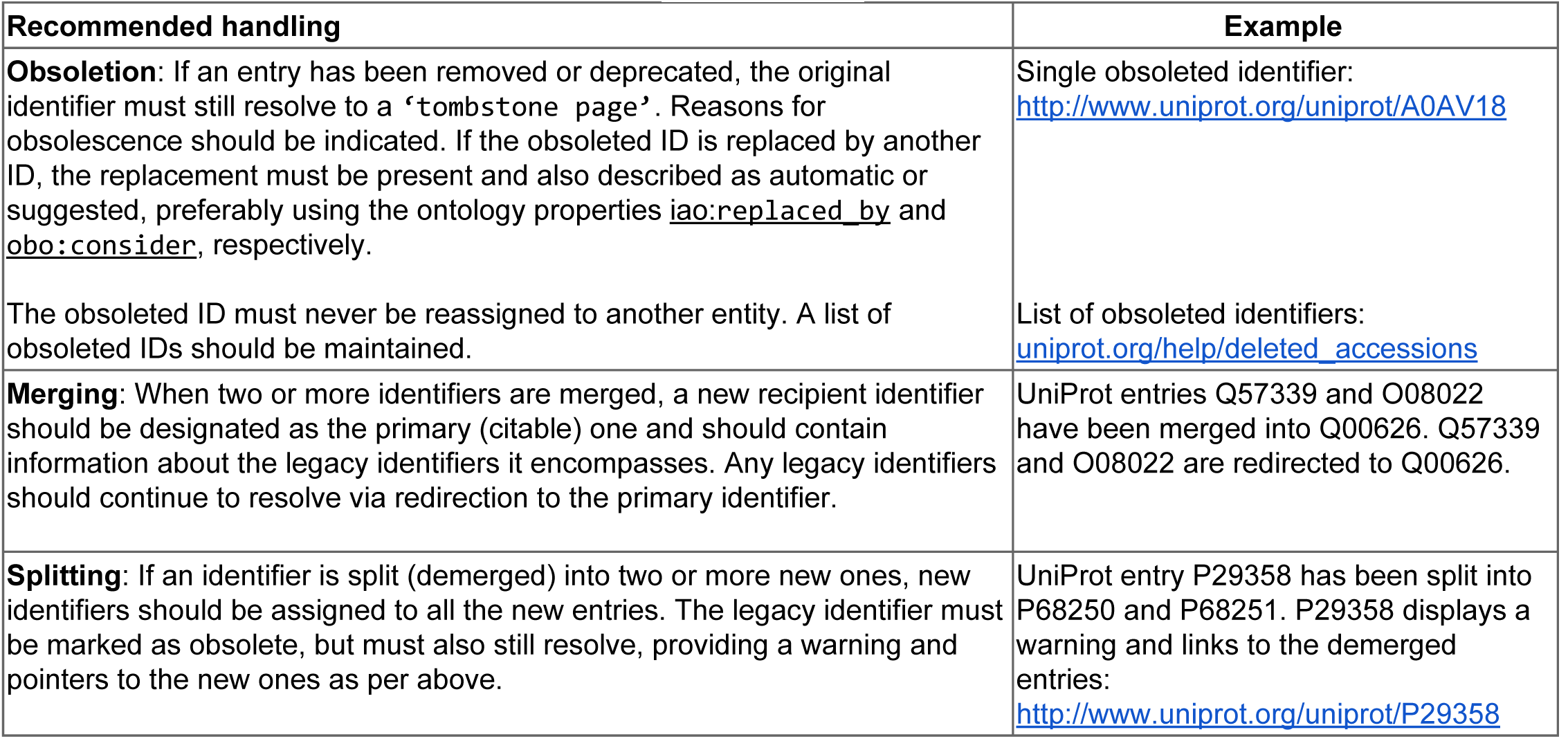
Recommendations for identifier lifecycle management.

### Lesson 8. Make URIs clear and findable

Persistent URIs almost always differ from the ephemeral URLs to which users are ultimately directed (Figure 2). Therefore, whether you produce original data, or reference others’ data, make persistent URIs obvious to users so that they are less inclined to ABC (Address Bar Copy). As a group, the best practitioners of this lesson are currently academic journals; they prominently advertise the DOI corresponding to each article. In situations where the version of a data record matters, advertise the corresponding “permanent link” (permalink) together with a statement about persistence. E.g.

> *“The permanent link to this page, which will not change with the next release of Ensembl is*: htto://e85.ensembl.ora/id/ENSMUSG00000033577 *We aim to maintain all archives for at least five years; some key releases may be maintained for longer”*

For archived records that are *out of date,* make this clear to the user and provide a link to the updated version (see http://www.uniprot.org/uniprot/Pi2345.1, for instance). Although it is good practice for each database website to include general citation guidance for users [26], it is increasingly important to provide a pre-populated citation *at the level of each record.* When it comes to making record-level citation clear on every page, eagle-i [27] provides the best example of a primary data source that we know of (outside of providers that issue DOIs) **(Figure 3)**. Additional features that are useful in such widgets are that full references should be copy-pastable, integrated with reference managers, and pre-populated with the version information and access date.

**Figure 3.**
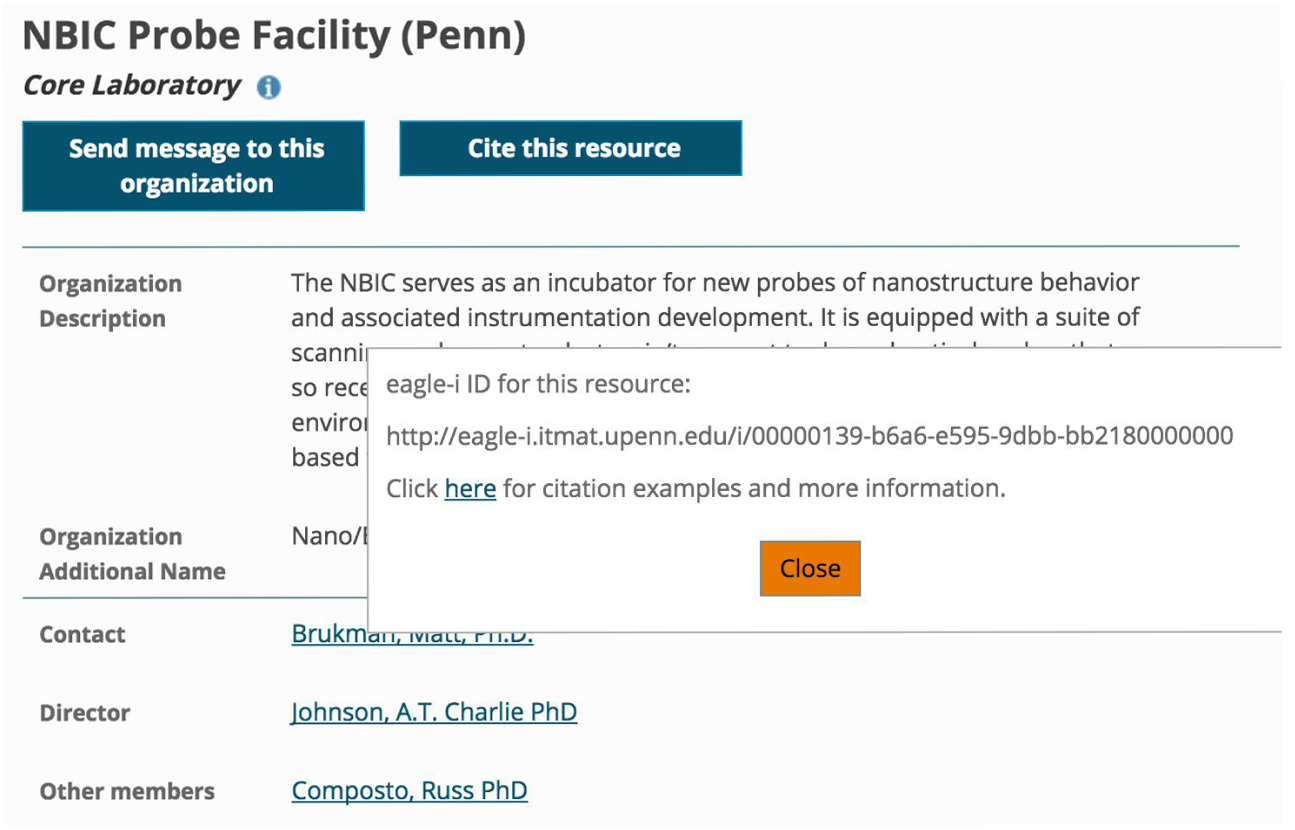
eagle-i record-level citation widget

### Lesson 9. Document the identifiers you issue and use

The global-scale identification cycle is a shared responsibility and provider/consumer roles often overlap in the context of data integration. Whether you issue your own identifiers or just reference those of others, you should document your identifier policies. **Supplemental Table S6** provides a set of questions that data providers and re-distributors can use to develop such documentation. Documentation should be published alongside and/or included together in a dataset description, for instance, as outlined in the recommendations for Dataset Descriptions developed by the W3C Semantic Web in the Health Care and Life Sciences Interest Group [28]. For examples of such documentation see ChEMBL [29] and Monarch [30]; the format may vary.

### Lesson 10. Reference and display responsibly

The final lesson describes referencing recommendations for data redistributors: data aggregators, who collect information from different sources and re-display it; data publishers, who disseminate scientific knowledge through publications; and online reference material such as WikiData [31].

When external entities are referenced in narrative online text, they should be hyperlinked to their URIs or to pages/metadata containing their URIs. Access URLs are volatile (see **Lesson 4**) and must not be used for referencing or linking in any context intended to persist.

Broader issues associated with citation of data and software in the traditional literature are outside of the scope of this paper, but **Text S1** lists relevant complementary efforts. Our recommendations regarding data citation in the literature are circumscribed: within static documents of record (eg. in PDFs), or in situations where link updates are costly/difficult, we strongly advocate always using the URLs of well-established third-party resolvers, whether they be primary resolvers such as doi.org or hdl.net or meta-resolvers such as identifiers.org, or n2t.net (Supplemental **Text S4**). Each provider has a corresponding URI pattern; however, those URIs can and do change over time. Third-party resolvers are not immune to change; the fact that the PURL.org resolver recently nearly sunset into “read-only” mode illustrates a) the importance of sustained community buy-in and governance and b) that reliance on 3rd parties for resolution is not without its risks. Nevertheless, the risk that URIs will break because of resolver change is modest and easier to mitigate compared to the risk that any *single* referenced collection will move or disappear. It is incumbent on meta-resolvers to be vigilant about detecting and updating their redirection rules in the face of provider changes. identifiers.org is able to redirect to one of a few potential provider destinations based on an algorithm that considers a) provider uptime, b) whether a given provider is a ‘primary’ source of the data in that collection. N2T.net and Identifiers.org recently joined forces [32] to harmonize identifiers in the same way, using the same prefixes. As part of this partnership, they have both have adopted simple syntax that gives users finer grained control, to request to be directed to a specific source of the data; for instance specifying the primary source of the data whether or not it has the best record of up-time.

Redistributors of data should monitor their references to other sources; any ‘dead’ links should be reported to the original data provider. If the original provider does not fix the broken link, your reference to it should be marked obsolete both visibly (for user interaction/interpretation), and within any accompanying metadata (for computational interaction/propagation). Differentiate identifiers linked internally within your application from identifiers linked outside your application; one way to do this is by using the linkout icon; consider opening all external links in a new browser window or tab in order to avoid confusion.

## Conclusion

Better identifier design, provisioning, documentation, and referencing can address many of the identifier problems encountered in the life science data cycle - leading to more efficient and effective science. However, it is well established that just because it is broadly agreed that a practice would be beneficial to the community, does not mean that it is adopted; to have an impact, the adoption of best practice has to be both easy *and* rewarding. In the broader context of scholarly publishing, this is just what DOIs afford; DOIs succeeded because they were well aligned with journals’ business goals (tracking citations) and because the cost was worth it to them. However, in the current world where everyone is a data provider, alignment with business goals is still being explored: meta resolvers can provide a use case for journals and websites seeking easier access to content, while software applications leverage these identifier links to mine for knowledge.

We recognize that improvements to the quality, diversity, and uptake of identifier tooling would lower barriers to adoption of the lessons presented here (**Text S7**). Those that issue data identifiers face different challenges than do those referencing data identifiers; we understand there are ecosystem-wide challenges that need will undertake to address these gaps in the relevant initiatives (**Text S1**). We also recognize the need for formal software-engineering specifications of identifier formats and/or alignment between existing specifications. Here, we implore all participants in the scholarly ecosystem - authors, data creators, data integrators, publishers, software developers, resolvers - to aid in the dream of identifier harmony and hope that this paper can catalyze such efforts.

## Acknowledgments

JA McMurry, T Burdett, N Juty, S Jupp, and C Morris were supported in part by the BioMedBridges project, which is funded by the European Union Seventh Framework Programme within Research Infrastructures of the FP7 Capacities Specific Programme, grant agreement number 284209. <u>EMBL-EBI core funds</u> supported H Parkinson, MJ Martin, J McEntyre, H Hermjakob, J Malone, M Courtot. <u>ELIXIR core funds</u> supported N Blomberg, R Jimenez. The European Commission provided additional support for Simon Jupp under grant number <u>601043</u> (“DIACHRON”) and for N Juty and H Hermjakob under grant number <u>312455</u> (“Infrastructure for Systems Biology - Europe (ISBE)”). The Drug Disease Model Resources grant number DDMoRe <u>115156</u> (“Innovative Medicines Initiative”) supported C. Laibe. Support was also received from the following BBSRC grants: <u>BB/L005050/1</u> (“ELIXIR-UK, Manchester”) for SA Sansone, A Gonzalez-Beltran and C Goble; <u>BB/M013189/1</u> (“DMMCore”) for C Goble, J Snoep, and N Stanford; <u>BB/K019783/1</u> (“Continued development of ChEBI”) and <u>BB/M006891/1</u> (“EMPATHY”) for N Swainston; <u>BB/M017702/1</u> (“SYNBIOCHEM”) for N Swainson and D Fellows; <u>BBS/E/B/000C0419</u> (“A systems approach to understanding lipid, Ca2^+^ and MAPK signalling networks”) for N Le Novère; <u>BB/L005069/1</u> (“ELIXIR-UK, Oxford”) for SA Sansone, A Gonzalez-Beltran and P Rocca-Serra. NIH support was provided from the following grants: <u>U41HG007822</u> (“UniProt”) for MJ Martin; <u>U24AI117966-01</u> (“bioCADDIE”) for SA Sansone, A Gonzalez-Beltran and P Rocca-Serra; <u>U54AI117925</u> (“CEDAR”) for M Dumontier, SA Sansone, A Gonzalez-Beltran and P Rocca-Serra; <u>R24OD011883</u> (“Monarch Initiative”) for CJ Mungall, MA Haendel, JA McMurry and NL Washington; <u>NHGRI P41HG002273-09</u> (“Gene Ontology Consortium”) for CJ Mungall. Additional support for CJ Mungall and NL Washington was received from the Director, Office of Science, Office of Basic Energy Sciences, of the U.S. Department of Energy under [Contract No. <u>DE-AC02-05CH11231</u>].

The authors wish to thank <u>Mary Todd Bergman</u>, <u>Ewan Birney</u>, <u>Fiona Cunningham</u>, <u>Richard Cyganiak</u>, <u>Adam Faulconbridge</u>, <u>Andrew M Jenkinson</u>, <u>Sirarat Sarntivijai</u>. <u>Stephanie Suhr</u>, <u>Eleanor Williams</u>, <u>Martin Fenner</u>, and <u>Tim Clark</u> for their valuable feedback and suggestions. We also wish to thank the BioMedBridges Scientific Advisory Board for the suggestion to address this important issue and the reviewers for their constructive comments.

## Supporting information

### Supplementary Text S1. Initiatives relevant to identifiers

- **BD2K (Big Data 2 Knowledge)** [33]. This US program supports a variety of initiatives aimed at making better use of the diversity of biomedical data, including various data integration efforts.
- **BioMedBridges** [34]. This is an implementation-driven project to integrate data that facilitates translational research [34].
- **DataCite** [35]: DataCite is interested in enabling the persistent identification of data, and develops and supports the standards required to achieve this [35].
- **DCIP** [36]: The Data Citation Implementation Pilot goal is to provide basic coordination between publishers, repositories and identifier / metadata services for early adopters of data citation according to the JDDCP [36].
- **Diachron** [37]: DIACHRON intends to address and cope with certain issues arising from the evolution of and identification of data in a web environment.
- **ELIXIR** [38]: A pan-European research infrastructure tasked with safeguarding and managing biological data.
- **Force11** [39]: This international pan-disciplinary organization is a forum for innovations in scholarly communication, including citation of data, research resources, and other web artifacts such as software.
- **Monarch Initiative** [40]: A global consortium dedicated to integrating cross-species genotype-phenotype data for disease discovery.
- **RDA** [41]: The Research Data Alliance is a globally active alliance interested in achieving the open sharing of data across countries, technologies and research domains.
- **W3C HCLS** [42]: The World Wide Web Healthcare and Life Sciences Interest group aims to develop semantic standards for interoperability.
- **OBO Foundry** [43]: The OBO Foundry consortium is a collaborative of ontology developers adhering to common best parctices and shared principles to ensure interoperability, including a common identifier and citation policy [44].
- **GA4GH** [45]: The members of the Global Alliance for Genomics and Health work towards integrating and analysing genomic data.
- **JATS** [46]: The Journal Article Tag Suite is an application of NISO Z39.96-2015, which defines a set of XML elements and attributes for tagging journal articles and describes three article models. JATS is a continuation of the NLM Archiving and Interchange DTD work begun in 2002 by NCBI [47]. It can also be used to cite data in journals.

**Supplementary Table S2.**
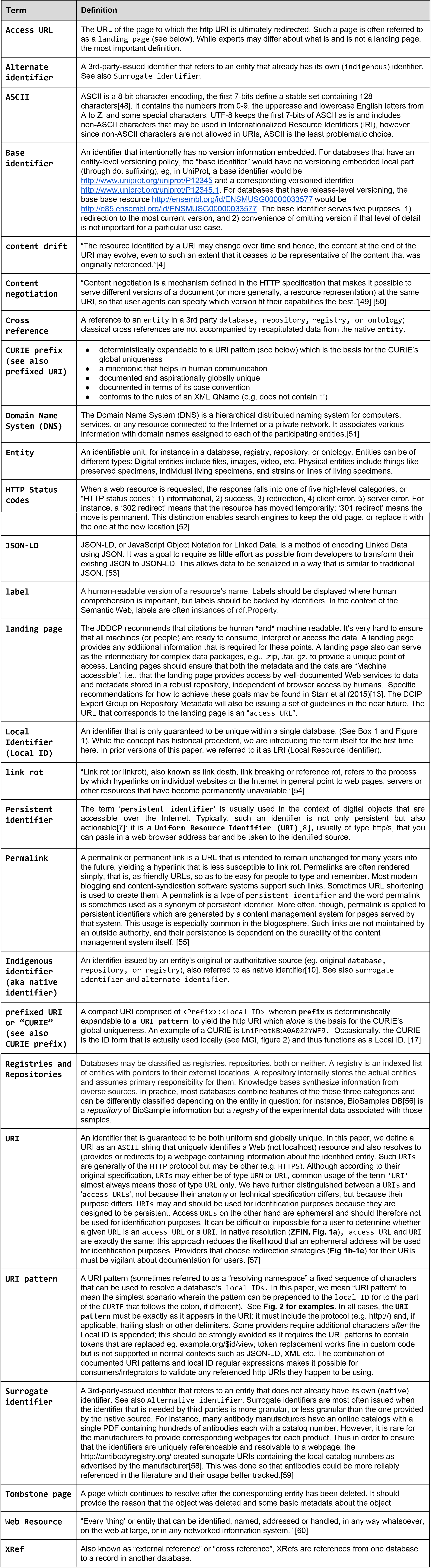
Glossary of web technology terms.

### Supplementary Text S3. Utility of CURIEs

The features that make for a good persistent URI also make for good CURIEs: desirable features include lack of semantics in both the URI pattern and the local ID (**Lesson 4**), absence of characters after the local ID (**Lesson 5**), omission of problematic characters etc (Lesson 5). CURIEs can complement http URIs in important ways for curators and data integrators:

A. **Brevity.** In the life sciences, prefixed identifier forms are traditionally favored over http URIs in curation tasks; for instance, within spreadsheets, online lab notebooks, and anywhere where identification is a core concern but where screen real estate is limited.
B. **Location-independence.** Third-party data integrators often add knowledge on top of existing identifiers, for instance as MonarchInitiative.org does with OMIM. But if Monarch’s URI’s instead included the embedded http URI of the OMIM source dataset it would look like https://monarchinitiative.org/uri=http://omim.org/entry/154700 instead of like https://monarchinitiative.org/OMIM:154700. If the OMIM ID were not converted to its CURIE form, the resulting URI in Monarch would be a) very long b) permanently vulnerable to any volatility in the original source URI. Encoding the prefix mappings for the sources dynamically provides both simplicity and
C. **Clues for collapsing equivalents.** Due to a lack of awareness and to evolving implementations and collection scope, it is exceptionally rare that only a single http URI is used for an entity. Although it is difficult to reliably ‘normalize’ equivalent URIs [61] that are syntactically different, the use of CURIEs can provide clues that facilitate it.

**Supplementary Table S4.**
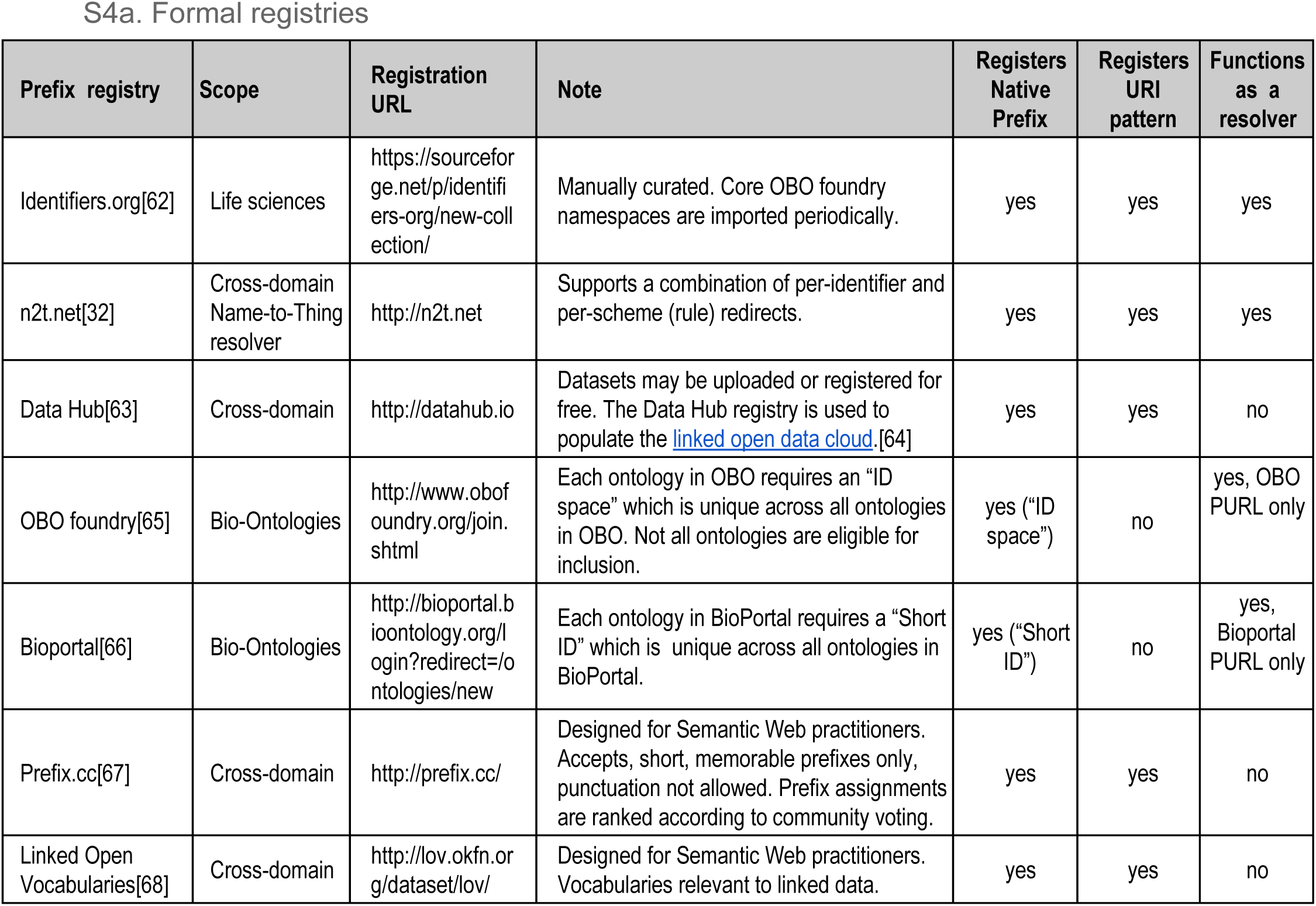
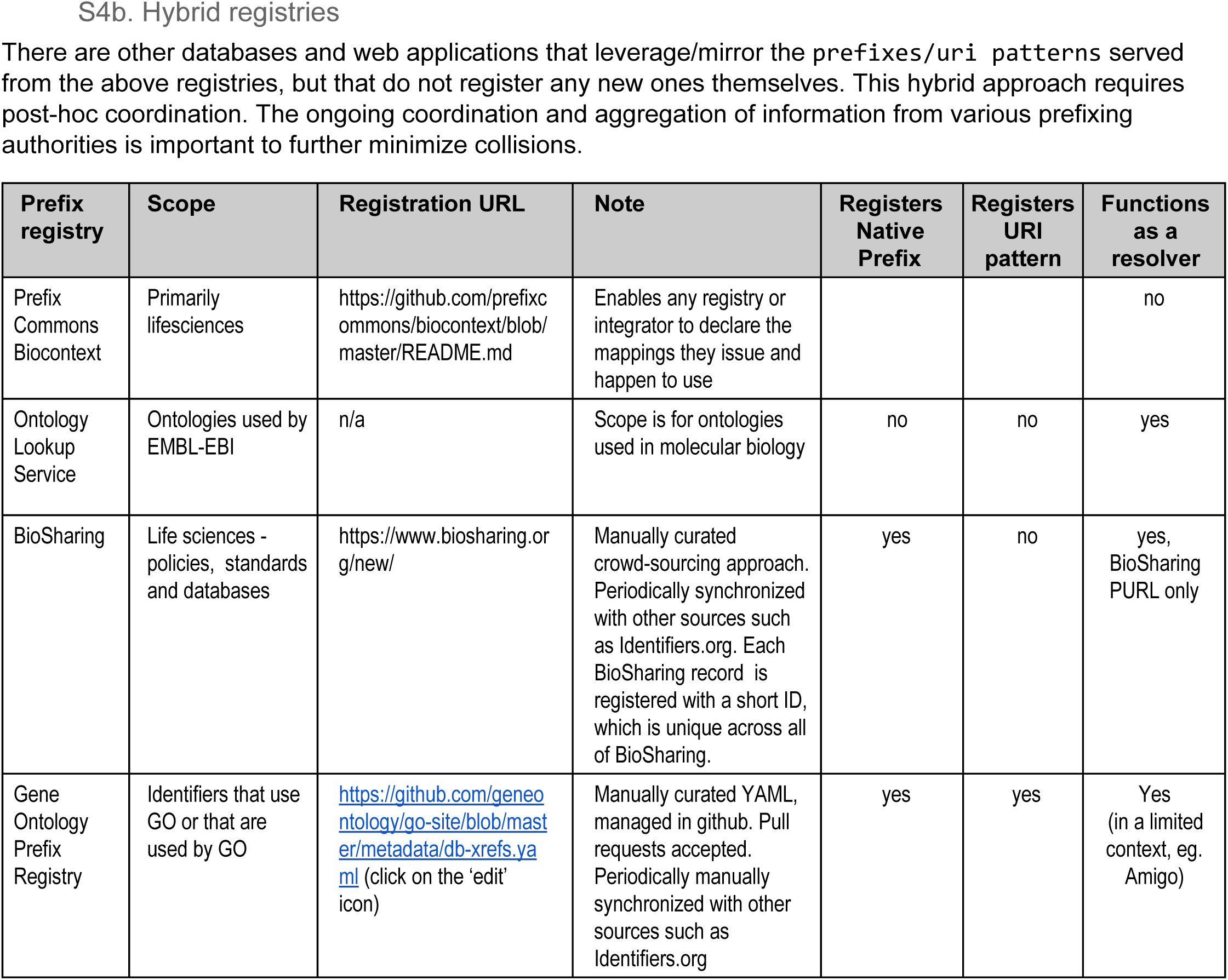
Prefix and URI pattern registries.

### Supplementary Text S5. Things to consider when choosing a resolver approach

There are basically three kinds of approaches to serving URIs on the web: (a) “native” URIs that require no redirection at all (as in Fig. 1, ZFIN). (b) “in house” URIs that redirect internally (as in Fig. 1, Ensembl); and (c) schemes using an external resolving authority (as in Fig. 1, Biosamples). Representative resolver authorities that meet the JDDCP (https://www.force11.org/datacitation, [69]) criteria are e.g. DOI (DataCite [35], CrossRef [70]), Identifiers.org [71], Handle.net [72], PURL [73], EPIC [74], N2T [75] and NBN [76]; these are described in Starr et al [13]. Additional resolver authorities that meet the criteria but which are not described therein are EPIC [77] and w3id [78]. Note that PURLs under the authority of PURL.org had gone into read-only mode and were therefore no longer adherent to the JDDCP principles; however, the InternetArchive [79] has assumed responsibility for them as of September 2016 [80].

Below are some additional criteria you may want to consider in choosing one of these resolvers.

- Does the resolver retain the native Local Resource Identifier that you issue (eg. identifiers.org, n2t.net), or does it instead issue a new one? (eg. DOI).
  ◦ If the resolver *does* issue a new identifier, what is the typical turnaround time between request and fulfilment? Can you obtain an identifier before you yourself need to use it?
- Would you or your institution need to pay fixed/variable costs to have your identifiers resolved? If the service is free for those that need their identifiers resolved, who pays to maintain the service?
- Is the service capable of issuing and managing identifiers in the kinds of volume you would require?
- Change management policy
  ◦ Will you need to change the data which is referenced by the URI, and if so does the resolving system under consideration permit such change?
  ◦ Is the object to which the URI resolves allowed to be removed?
  ◦ Does the resolver support numerical suffixing for versions of the LRI?
  ◦ If new LRIs are issued for each version of an entity, how can versions be related to each other?
- Will you require the resolver to support multiple resolving locations (mirrors)?
- Does the resolver support content negotiation at resolver’s HTTP URI?
- Does the resolver collect, index, and/or curate metadata about individual entities?
  ◦ If so, is the metadata that is collected relevant for the types of entities identified?
- Does the resolver collect, index, and/or curate metadata about collections of entities (e.g. whole databases)?
  ◦ If so, is the metadata that is collected relevant for the type of collection?
- Does the resolver support controlled access for confidential data?
- Is the resolver cross-discipline?

**Supplementary Table S6.**
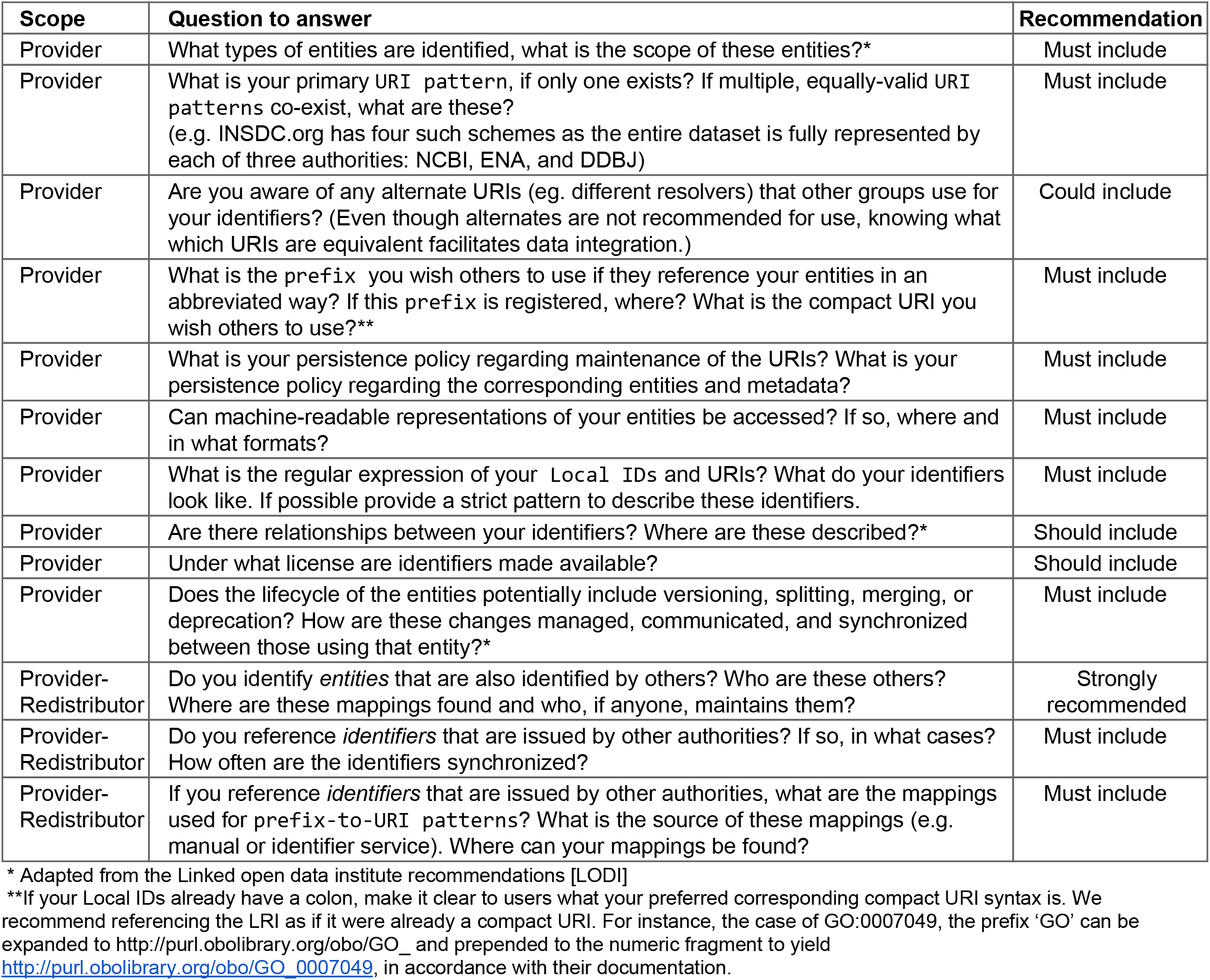
Questions that good identifier documentation should answer.

### Supplementary Text S7. Current and future efforts that would help lower barriers to adoption

#### Current efforts

- **Registries, 3rd party resolvers:** A list of identifier resolvers and identifier registries is in **Supplemental Table S3**.
- **PICR** [81]: Protein Identifier Cross-Reference Service has a service that returns identifier mappings, optionally including deleted ones. PICR or a similar service could be developed to have broader scope.
- **HCLS** [28]: Health Care and Life Sciences dataset descriptions provide a standard representation of the original sources of data (and therefore identifiers) in any integrated dataset.
- **JATS** [82]: In the context of the literature, Journal Article Tag Suite provides a standard way for data citations to be represented in the literature, facilitating credit and reward mechanisms. However, outside of the literature, referencing and display is primarily an issue of increasing awareness.
- **BioSchemas.org** [83] is promoting more consistent adoption of schema.org markup in the life sciences. Markup can facilitate more transparent provenance and credit mechanisms of integrated data, as well as optimizing data for discovery by search engines, whether Google, or others.

#### Future efforts

- **Identifier validator:** Identifier designers could help data producers choose the design that best suits their particular use case, validators could determine whether an existing identifier is valid according to a published scheme.
- **Embeddable citation widgets or citation markup** could help providers display citation information, clearly and consistently.
- **Archiving services: For archival of content, client-facing services include the Memento web protocol** [2]. We authors of this paper are not aware of any existing platforms that providers can outsource their content archiving to, but such a service may be worthwhile. Another function for archival services is for maintaining a robust network of linked entities. In this case, full archival of content may not be needed. Rather, resolver and/or indexing services may provide “tombstone pages” with essential metadata so that these entities can still be resolved.

